# A novel network controllability algorithm to target personalized driver genes for discovering combinational drugs of individual cancer patient

**DOI:** 10.1101/571620

**Authors:** Wei-Feng Guo, Shao-Wu Zhang, Tao Zeng, Luonan Chen

## Abstract

Treating cancer in precision medicine, it is important to identify the personalized combinational drugs under consideration of the individual heterogeneity. Many bioinformatics tools for the personalized driver genes identification have presented promising clues in determining candidate personalized drug targets for the personalized drugs discovery. However, it has not been studied how to fill the gap between personalized driver genes identification and personalized combinational drugs discovery. In this work, we developed a novel algorithm of structure network Controllability-based Personalized driver Genes and combinational Drug identification (CPGD), aiming to mine the personalized driver genes and identify the combinational drugs of an individual cancer patient. On two benchmark cancer datasets, the performance of CPGD for predicting the clinical efficacious combinational drugs is superior to that of other state-of-the-art driver gene-focus algorithms in terms of precision accuracy. In particular, by quantifying and referring the relationships between target genes of pairwise combinatorial drugs and disease module genes on breast cancer data set, CPGD can significantly divide patients into the discriminative high-risk and low-risk groups for risk asessment in combination therapy. In addition, CPGD can further enhance cancer subtyping by providing computationally personalized side effect signatures for individual patients. Collectively, CPGD provided a new and effecient bioinformatics tool from structure network controllability perspective for discovering personalized combinational drugs with personalized side effect consideration, so as to effectively support personalized risk assessement and disease subtyping.

**Significance:** It is quite challenging to predict personalized combinational drugs rather than patient-cohort‘s drugs based on cancer omics data. In this work, a novel structure network Controllability-based algorithm (CPGD) from feedback vertex sets control perspective was developed, for discovering efficacious combinational drugs of an individual cancer patient by targeting the personalized driver genes. The CPGD contains three methodological advances by exploring more precise mathematical models on high-throughput personalized multi-omics data. The first is that a proper network structure is constructed to characterize the gene regulatory mechanism of an individual patient. The second is that considering the weight information of network edges/relations improves the performance for predicting clinical efficacious combinational drugs compared with other drivers-focus methods. And the third is that proper evaluation metrics for personalized combinational drugs prioritization, personalized risk assessment and disease subtyping are designed when evaluating the performance of CPGD.

## Introduction

It is difficult to achieve the desired clinical effect for monotherapy and the usage of multiple drugs have demonstrated great advantages in overcoming drug resistance and improving clinical outcomes in disease therapy^1^. Consequently, combination therapy has been widely used in the treatment of cancer^2^. Furthermore, researchers have recognized that cancer is a heterogeneous disease, because tumor genes cooperate as well as adapt and evolve to the changing conditions for individual patients^3–6^. Thus, it is essential to consider individual heterogeneity for combination therapy in the treatment of cancer. However, the number of potential combinatorial drugs is astronomical, and these potential compound combinations cannot all be validated in a rational and rigorous manner for individual patients. Therefore, it is quite challenging to predict personalized combinatorial drugs rather than patient-cohort‘s drugs.

Increasing evidences show that cancer is induced by multi-gene genetics and environmental factors, rather than by a single target^7^, thus drug development is shifting from single-target based “one drug one target” paradigms to multitarget based combination paradigms in drug discovery^8^. Due to the rapid development of high-throughput screening, the emergence of system biology or network biology have raised the possibility of exploring multi-target intervention methods with synergistic effects for cancer treatment^9,10^. It is known that designing novel combinatorial strategies to target fundamental cancer driver genes (i.e., candidate drug targets) may result in synergistic antitumor activity and broader application of current therapies^11^. Some methods have been developed for driver gene identification with multi-dimensional genomic data, such as DriverML^12^, DriverNet^13^, MutSigCV^14^, OncoDriveFM^15^, SCS^16^ and DawnRank^17^. But, the goal of discovering personalized combinatorial drugs is far from being achieved, and new models and algorithms are still urgently required to identify the personalized driver genes for recommending the personalized combinatorial drugs in cancer treatment.

In this challenging field,structure network controllability methods have already offered a promising framework to identify cancer driver genes with novel biological and biomdical insights, helping us determine the candidate drug targets for drug discovery^18^. Structure network controllability methods mainly focus on how to identify driver nodes that are imposed by the control signal for driving the entire network toward a desired state^19^. In the past decade, many researchers have studied the structure network controllability principles, such as maximum matching set (MMS)^19^, minimum dominating set (MDS) ^20^, and feedback vertex set (FVS)^21,22^ Although these methodological advances have raised the possibility of exploring more precise mathematical models on high-throughput personalized multi-omic data for the discovery of efficacious personalized drug combination therapies, the existing structure network controllability methods face three limitations. The first is that a proper network structure is not available to characterize the gene regulatory mechanism of an individual patient, which is a rate-limiting step of structure network controllability methods. The second is that overlooking the weight information of network edges/relations may generate multiple driver-node sets resulting in a potential bottleneck for identifying the combinational drugs of the individual patients. And the third is that gold standard evaluation metrics are not available when evaluating the performance of identifying personalized combinational drugs by different structure network controllability methods. To overcome above limitations, we developed a novel network controllability-based algorithm (namely CPGD) to detect personalized driver genes and further identified the personalized combinational drugs of an individual patient. We first used a paired single sample-network method (Paired-SSN) to construct the personalized gene interaction network in which the interactions determine the state transition of an individual patient during cancer development. According to feedback vertex sets (FVS)-based controllability perspective, we then developed a novel nonlinear controllability method (namely Weight-NCUA) to identify the personalized driver genes for determining the state transition of the individual biological systems between disease state and normal state. In contrast with other structure-based network controllability approaches, Weight-NCUA considers the edge weight information (i.e., network scores) for the driver node optimizations. Finally, the information of sub-networks between the drugs and personalized driver genes was used for: i) prioritizing personalized combinatorial drugs to evaluate the ability of predicting clinical efficacious combinational drugs, ii) exploring risk assessment of paiwise combinatorial drugs, and iii) enhancing cancer subtyping by side effect quantification on personalized driver genes.

We evaluated the effectiveness of CPGD on the datasets of the breast invasive carcinoma (BRCA) and lung squamous cell carcinoma (LUSC) which are derived from The Cancer Genome Atlas (TCGA). These results show that: i) Compared with other state-of-the-art methods, CPGD can more effectively predict clinical efficacious combinational drugs. Furthermore, the prior gene interaction network can improve the prediction accuracy of combinational drugs. ii) CPGD detected 11 drug pairs which can actually divide patients into discriminative high-risk and low-risk groups and 8 efficacious drug pairs whose combination therapy on risk assessment are indeed better than that of a single drug treatment, by exploring the relationships between target genes of pairwise combinatorial drugs and disease module genes for BRCA cancer dataset. By quantifying the side effect of personalized combinatorial drugs on personalized driver genes for each individual patient, CPGD identified 3 cancer subtypes on breast cancer dataset. Of note, the greater the side effects of a specific patient subtype, the higher the fraction of patient deaths, and the higher the fraction of patients within later clincial stages (e.g., stage III and stage IV). In conclusion, we provided a new and effecient bioinformatics tool from structure network controllability perspective for discovering personalized combinational drugs with personalized side effect consideration.

## Results

### Algorithm overview of CPGD

From dynamical system viewpoint, gene expressions of an individual patient are the biological system variables, varying at different time points. It is the personalized gene interaction network (i.e., system structure) that results in the change of gene expression value (i.e., system variable)^23,24^ Therefore, CPGD hold a key assumption that the personalized gene interaction network determines the state transition between normal state and disease state of an individual cancer patient. The input of CPGD is the gene expression data of paired samples (i.e., normal and tumor samples) for individual cancer patient. Meanwhile, the main outputs of CPGD include: i) prioritization of personalized combinatorial drugs; ii) risk assessment for individual patients dependent on personalized drug pairs; iii) cancer subtyping by side effect quantification of personalized drug pairs on personalized driver genes. Due to the sufficiently avalable drug information of BRCA and LUSC, we collected their gene expression data from TCGA in this work. BRCA and LUSC datasets consist of 112 and 49 paired samples (i.e., normal and tumor samples for each patient), respectively.

As shown in **Figure 1**, CPGD mainly comprises the following three steps as follows:

**Step 1 Constructing the personalized gene interaction networks with Paired-SSN method**. CPGD used the method of Paired Single-Sample-Network (Paired-SSN) ^25^ to construct the personalized gene interaction networks for individual patient (**Materials and Methods**). The personalized gene interaction networks are constructed to reveal the significant gene interactions between the normal and tumor samples for each individual patient during cancer development. In this work, the personalized gene interaction networks are weight graphs, in which nodes represent genes, and edges denote the significant co-expression difference of gene interaction between the normal state and tumor state. The edge weights indicate the differential fold change of co-expression (strength) between normal sample and tumor sample for an individual patient.
**Step 2 Identifying personalized driver genes with Weight-NCUA method**. In this work, we are interested in how to determine the state transition of the gene expression level between disease state and normal state with the personalized driver genes. Thus, based on Feedback Vertex Set, we introduced a novel structural network controllability method (namely Weight-NCUA) to identify the personalized driver genes. The personalized driver genes are thought to be the candidate drug targets which can drive individual biological system from disease state to normal state (or approximate normal state) through drug activation signals. By using the weight information of nodes in the personalized gene interaction network, Weight-NCUA optimizes the personalized driver genes. Weight-NCUA tries to design a fitness index for representing the impact quality of personalized driver gene set, and to determine the personalized driver genes by identifying optimal FVS with maximum impact quality in the personalized gene interaction network.
**Step 3 Screening the role of personalized combinatorial drugs by tageting the personalized driver genes**. The role of personalized combinatorial drugs is screened by using the following aspects: i) Prioritizing the potential personalized anti-cancer combinatorial drugs by measuring the number of targeting personalized driver genes for a given drug combination in the interaction network of combinatorial drugs and genes. ii) Exploring risk assessment of paiwise combinatorial drugs. That is, we first selected top 10 candidate combinational drugs for each patient. Among these candidate combinational drugs, we selected the drug pairs with Complementary Exposure (e.g., drug pairs can both hit the personalized driver genes overlapping with disease module but have non-overlapping targeted personalized driver genes). Then we chose the efficacious drug pairs with signifacnt survival analysis results (p-value<0.05) by using their targeting personalized driver genes as the input of *SurvExpress* tool^26^ for risk assessment. iii) Enhancing the cancer subtyping with side effect quantification on personalized driver gene. That is, we calculated the *side effect score* (**Materials and Methods**) of each drug pair by quantificating side effect on personalized driver genes in the drug-target network with activation and inhibition interactions for each patient. Based on the *side effect score* of each drug pair, we obtained the number of drug pairs with an *aggravating effect* (*side effect* score >0), and the number of drug pairs with *enhancing effect* (side effect score <0), which are two features of side effect signatures of individual patients.

**Fig.1.**
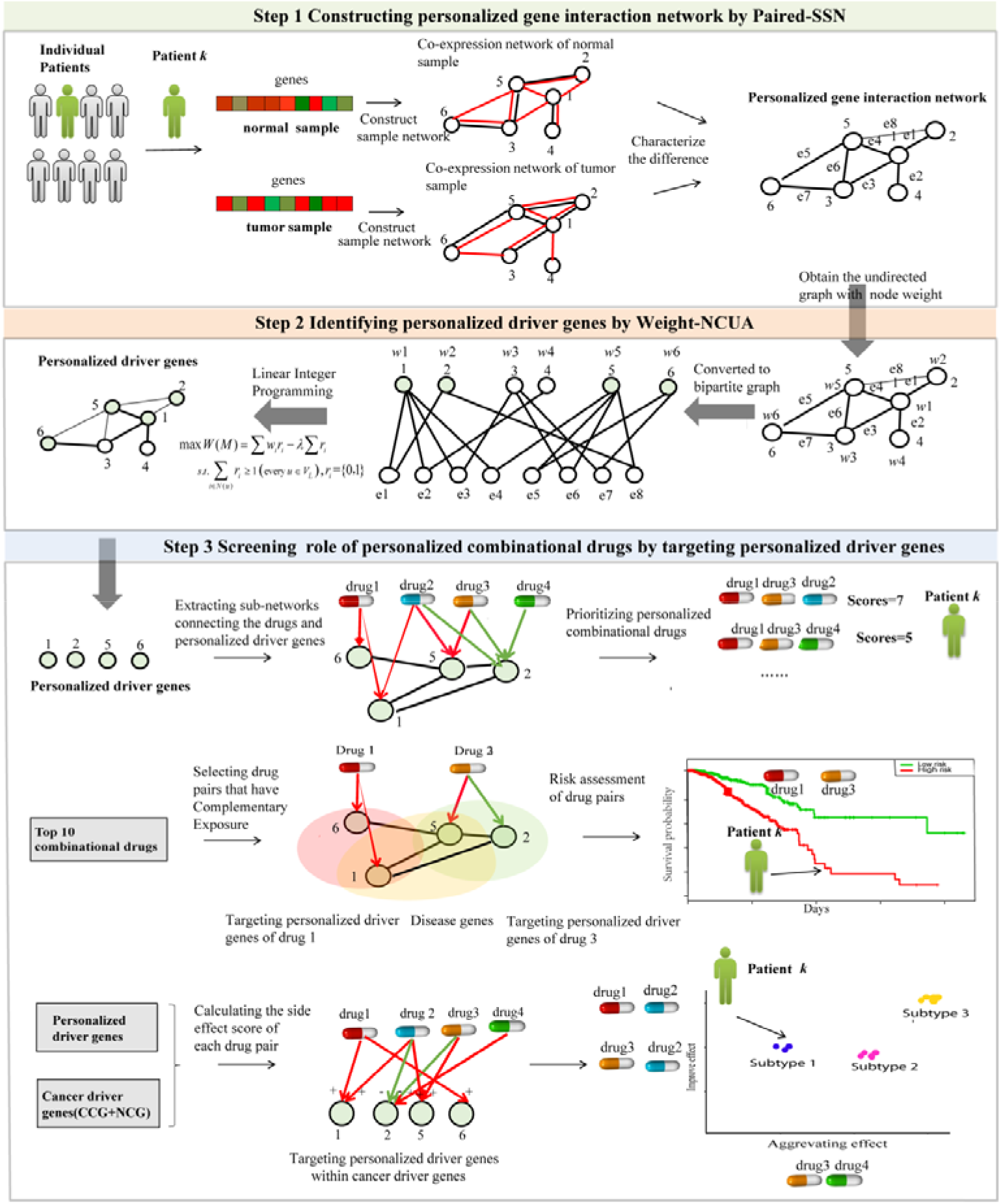
CPGD overview. First, a paired single sample network method (Paired-SSN) is used to construct the personalized gene interaction networks for capturing the phenotypic transitions between normal state and disease state. The personalized gene interaction networks are weight undirected graphs in which edges denote the significant co-expression difference between normal state and tumor state in the gene interaction networks, and edge weights indicate the differential fold change of co-expression between normal sample and tumor sample for a specific patient. Then, a structure network controllability method (Weight-NCUA) is developed to identify the personalized driver genes, where the driver genes are considered as candidate drug targets towards the desired control objective by drugs activations. Finally, we screened the role of personalized combinatorial drugs from the following aspects: i) Prioritizing the personalized combinatorial drugs by measuring the number of targeting personalized driver genes in the interaction network between combinatorial drugs and genes. ii) Exploring risk assessment of drug pairs. That is, we selected drug pairs with Complementary Exposure from top 10 candidate combinational drugs for each patient as candidate, then selected efficacious drug pairs with significant survival analysis results (p-value<0.05) for risk assessment. iii) Enhancing cancer sub-typing by side effect quantification on personalized driver genes. When two drugs act simultaneously on the same target, their action of two combinations (+, +) and (–, –) will be referred to as *coherent* action, and the action of two combinations (+, –) is called incoherent action. By attaching signs to the mechanisms of action, the *side effect score* can be calculated in the drug-target network with activation and inhibition interactions for each patient. Based on the side effect score of each drug pair, we can obtain the number of drug pairs with aggravating effect (*side effect score* >0), and the number of drug pairs with enhancing effect (*side effect score* <0) as two features of side effect signatures of individual patients.

### Determination of the gene interaction networks and parameters in CPGD

To assess the effect of different sources of gene interaction networks on the performence of CPGD, we adopted three gene interaction networks (i.e., Network1, Network2 and Network3) from different literatures. The reference Network1 is built by Hou *et al*.^17^, which consists of 11,648 genes and 211,794 edges by integrating a variety of data sources, such as MEMo^27,28^, Reactome^29^, NCI-Nature Curated PID^30^, and KEGG^31^. The reference Network2 is built by Quan *et al*^32^ from the Synthetic Lethality genes interactions Database (SynLethDB), which consists of 6,513genes and 19,955 synthetic lethal gene pairs for human tumors. The reference Network3 is constructed by Vinayagam *et al*. ^33^, which consists of 6,339 proteins and 34,813 edges, where the edge denotes the hierarchy of signal flow between the interacting proteins.

By using each of three reference gene interaction networks, CPGD can output personalized driver genes for each individual patient, and rank the candidate combinatorial drugs acorrding to the number of targeting personalized driver genes. Based on the ranking of candidate combinatorial drugs for each individual patient, we assess the effect of different reference gene interaction networks on CPGD by using the following metric.

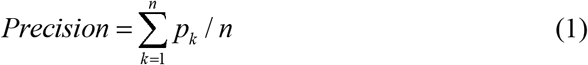

where *p_k_* denotes the fraction of the top *k* predicted combinatorial drugs within the Clinical Anti-cancer Combinational drugs for treating cancer, and *n* is the number of top ranked anti-cancer drug (here, *n* =10).

The performance of CPGD with different reference gene interaction networks on BRCA and LUSC datasets are shown in **Figure 2A and Figure 2B**. According to the results in figure 2A and 2B, we choose the Network2 with the highest *Precision* as the reference gene interaction network for BRCA, and Network1 with the highest *Precision* as the reference gene interaction network for LUSC. Besides, the balance parameter (*λ*) has different effects for CPGD on BRCA and LUSC cancer datasets and the reference networks. From Figure 2A and 2B, we can see that the precision of CPGD with *λ* = 100 is the highest. Thus, we selected *λ* = 100 for BRCA and LUSC in this work.

**Fig.2.**
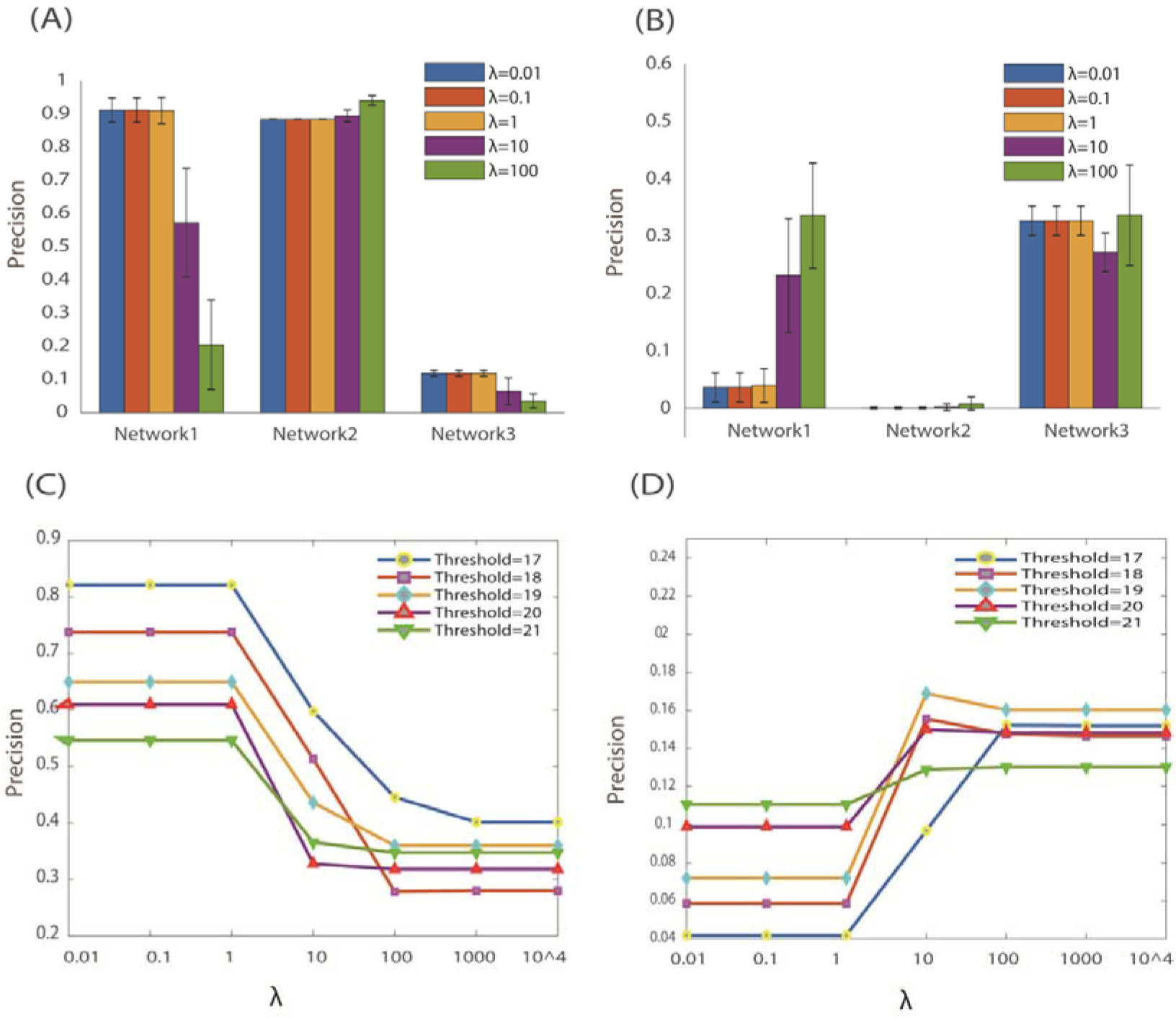
Effect of the reference gene interaction networks and parameters in CPGD. (**A**) Results of balance parameter (*λ*) on Network1, Network2 and Network3 for BRCA. (**B**) Results of balance parameter (*λ*) on Network1, Network2 and Network3 for LUSC. (**C**) Results of only considering the gene expression data without the gene interaction data for BRCA. (D) Results of only considering the gene expression data without the gene interaction data for LUSC. Since there are too many edges in the gene co-expression network, we separately chose the threshold of 17, 18, 19, 20 and 21 to filter the edges with low weight scores which are calculated with formula 4.

To further demonstrate the importance of prior-known gene interaction networks, we recalculate the *Precision* of the predicted combinatorial drugs by directly utilizing the expression network alone. The Precision of CPGD on the expression network with different thresholds (**Figure 2C and 2D**) is less than that on the prior-known gene interaction networks (**Figure 2A and 2B**), demonstrating that a map of prior-known gene interaction networks indeed help improve the discovery of efficacious combinatorial drugs. In one word, the choice of proper prior-known network structure should be an important factor for CPGD.

### Comparisions and evaluation of CPGD with other existing methods

The key contribution of our CPGD is to identify driver genes for inferring combinatorial drugs.To evaluate the effectiveness of CPGD, we compare our CPGD with other state-of-the-art driver genes identification methods, such as DriverML^12^,DriverNet^13^, MutSigCV^14^,OncoDriveFM^15^, SCS ^16^, DawnRank^17^and ActiveDriver^34^. In addition, we also compare CPGD with the traditional differential expression gene (DEG) approaches, i.e. DEG-FoldChange, DEG-p-value and DEG-FDR. DEG-FoldChange method selects the personalized driver genes by calculating the fold-change (i.e., |log2(fold-change)|>1) between the normal sample and tumor sample of an individual cancer patient. DEG-p-value and DEG-FDR methods select the personalized driver genes by respectively calculating the p-value and FDR (<0.05) between a tumor sample for an individual cancer patient and a group of control samples collected from the individual patients.

The results of CPGD and other methods are shown in Figure 3, from which we can see that the *Precision* of CPGD is higher than that of other 8 driver gene identification methods and 3 DEG identification methods on two benchmark cancer datasets. These results indicate that the ability of our CPGD for predicting clinical efficacious combinational drugs is superior to other driver gene-focus methods.

**Fig.3.**
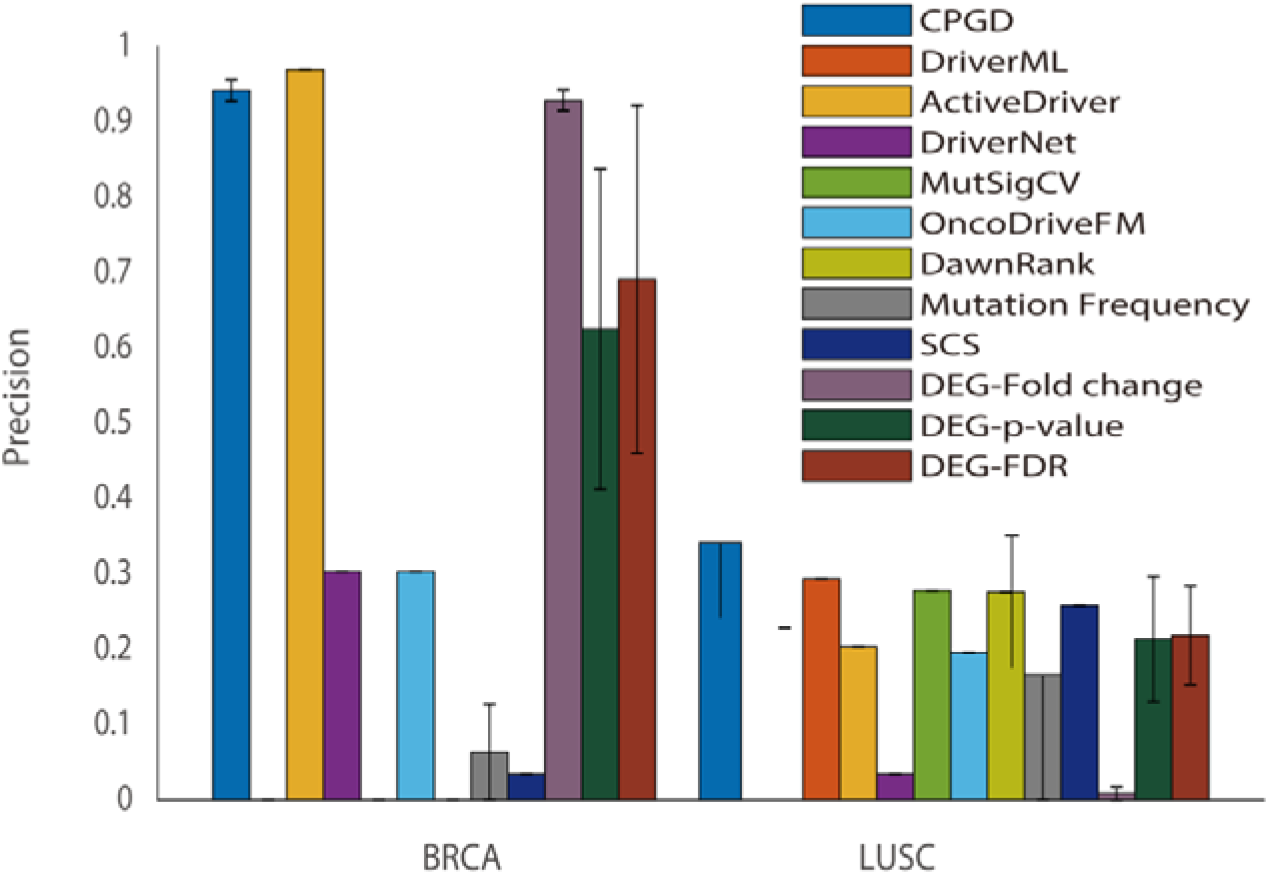
The comparison and evaluation of CPGD and other existing methods for predicting clinical efficacious combinational drugs on BRCA and LUSC cancer datasets. The y-axis denotes the *Precision* of the top 10 predicted combinatorial drugs. The bar colors represent different algorithms.

### Influence of single sample network construction methods

In order to investigate the effect of different network construction methods on CPGD, we separately use the methods of Paired-SSN^25^, CSN^24^, SPCC^35,36^ and LIONESS^37^ to construct the personalized gene interaction networks. For SPCC and LIONESS methods, after obtaining the SPCC and LIONESS co-expression distribution (*S*) of all gene pairs, we choose a threshold *W* to filter the edges with low co-expression value.

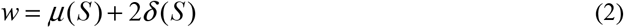

where *μ*(*S*) and *δ*(*S*) are the mean and standard variance of *S* of all gene pairs. Similarity with Paired-SSN^25^, CSN^24^, SPCC^35,36^ and LIONESS^37^ methods, we separately choose the Network2 and Network1 as the reference gene interaction network for BRCA and LUSC.

The results of our CPGD by using four single sample network construction methods are shown in Figure 4, from which we can see that the precision of Paired-SSN and CSN is higher than that of SPCC and LIONESS methods, and the precision of Paired-SSN and CSN is almost equal. In this work, we select the Paired-SSN method to construct the personalized gene interaction network.

**Fig.4.**
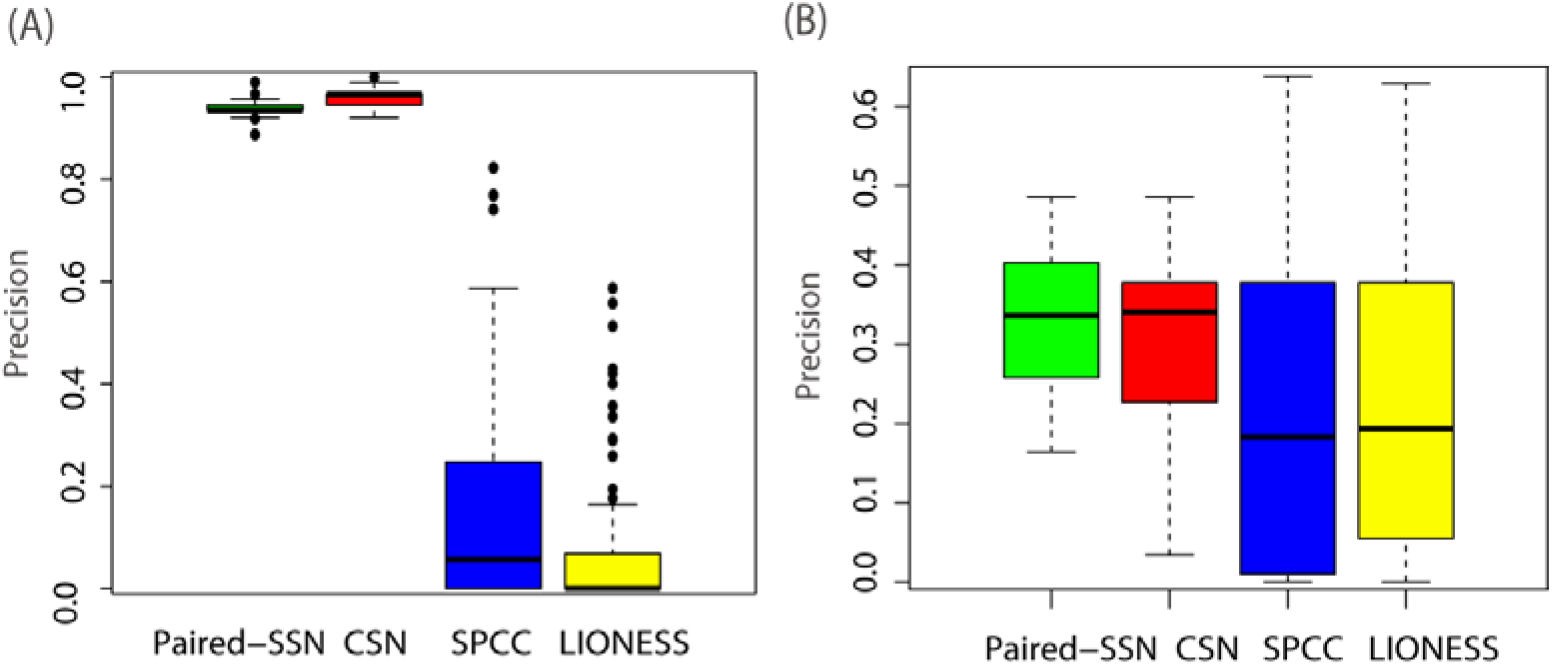
Influence of different single sample network construction methods on CPGD. To evaluate the usage efficiency of single sample network construction methods (i.e., Paired-SSN, CSN,SPCC and LIONESS) for personalized drug discovery, the combinatorial drugs annotated in the Clinical Anti-cancer Combinational drugs (CAC) are applied to obtain the *Precision* of the top-ranked/predicted anti-cancer combinatorial drugs for BRCA (A) and LUSC (B).

### Influence of network controllability methods

A key aspect of CPGD is to use the metwork controllability method of Weight-NCUA to identify the personalized driver genes. To evaluate the influence of different network controllability methods for CPGD, we compare Weight-NCUA with other structure network controllability methods, such as MMS^19^, MDS^20^ and DFVS ^21,22^on the personalized gene interaction networks built with Paired-SSN^25^ for predicting clinical efficacious combinational drugs (**Figure 5**). As shown in **Figure 5**, among the top 10 predicted combinatorial drugs on BRCA and LUSC, the *Precision* of Weight-NCUA respectively arrives 92.39% and 37.97%, which are higher than that of other structure network controllability methods. The main reason is that Weight-NCUA considers the network scores for the optimization of driver nodes. Figure S2 in **Additional file 1** shows that the number of driver nodes for Weight-NCUA is not minimum among all the structure network controllability methods, indicating that minimum number of driver nodes is not the unique objective for controlling the dynamic of the network. Thus, the performance of Weight-NCUA is better for identifying driver genes and recommending efficacious combinatorial drugs due to its conisderation in biolgocial and biomedical context.

**Fig.5.**
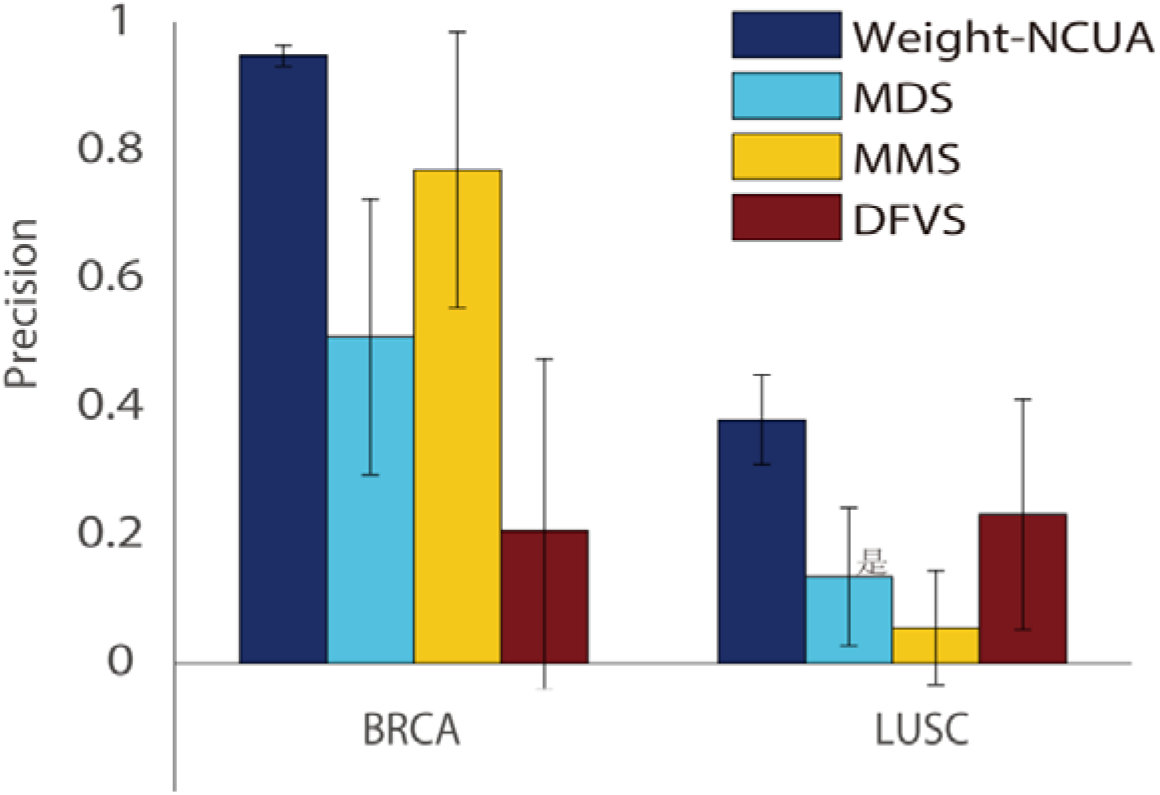
Results of Weight-NCUA, MDS, MMS and DFVS network controllability methods for top-ranked/predicted anti-cancer combinatorial drugs on BRCA and LUSC, respectively.

### Structural and functional property of personalized driver genes

After determing the suitable reference newtork source, single sample network construction method and network controllability method, CPGD can effectively identified a group of personalized driver genes. For analyzing the functional and structural properties of the personalized driver genes, we implement the mutation profile test and epistasis-interaction test (**Materials and methods**) on the BRCA and LUSC. The mutation profile test evaluates whether the personalized driver genes are significantly enriched in the mutated genes for an individual patient. The mutated genes of BRCA and LUSC are obtained from the single nucleotide variations (SNVs) data on TCGA ^38^ database. The epistasis-interaction test evaluates whether the personalized driver genes are epistatic, that is, comparing with random gene pairs, the personalized driver genes are significantly connected in the gene interaction network. The results of mutation profile test and epistasis-interaction test on BRCA and LUSC are shown on **Figure 6**. From **Figure 6**, we can find that: i) The P-values of most patients in mutational level test for BRCA and LUSC are less than 0.05 (ESg =-log_10_(p-value)<2), indicating that personalized driver genes tend to be the mutated genes in the mutational profiles of most patients. ii)The P-values in the epistasis-interaction test for BRCA and LUSC are less than 0.05 (ESg=-log_10_(p-value)<2), suggesting that the personalized driver genes are significantly and functionally connected in the gene interaction network of cancer patients. The results of the mutation profile test and epistasis-interaction test show that the personalized driver genes can be integratively characterized on mutational profile and network level.

**Fig.6.**
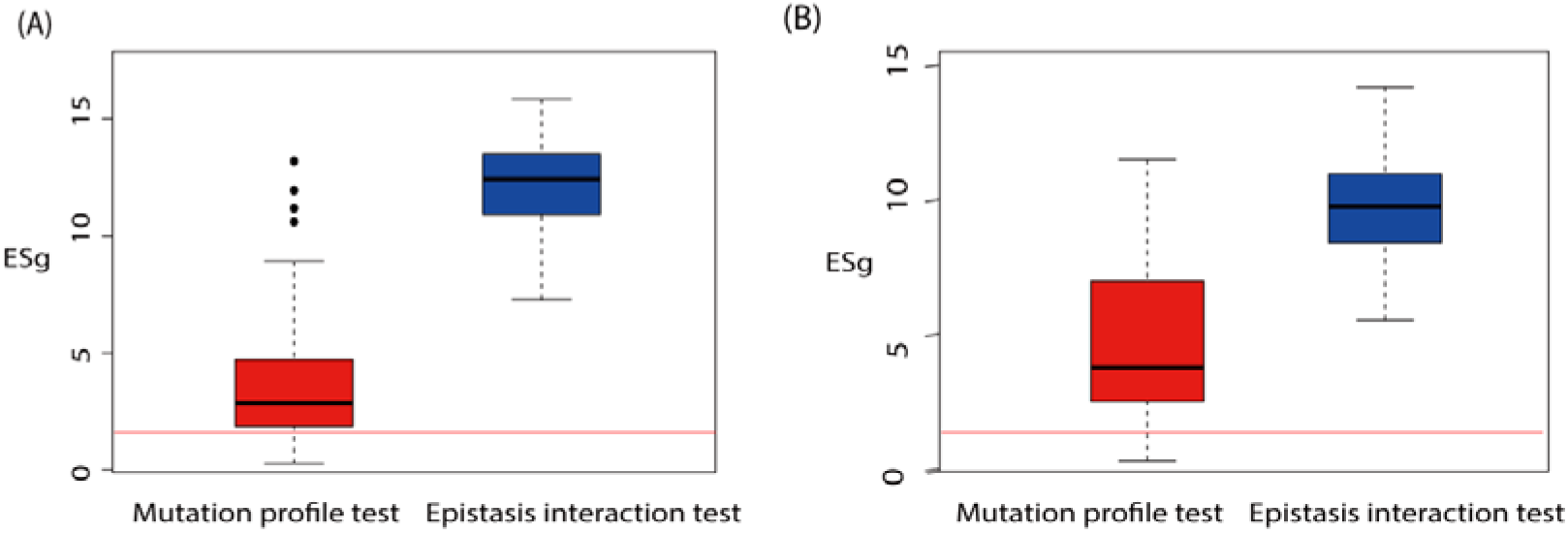
Functional and structural property of personalized driver genes. (A) Results of the mutational level, and epistasis-interaction level (or network level) tests for personalized driver genes identified with CPGD on BRCA. (**B**) Results of the mutational level, and epistasis-interaction level (or network level) tests for personalized driver genes identified with CPGD on LUSC. The Enrichment Scores of driver genes is defined as ESg=-log_10_(p-value). The red line denotes ESg=2.

To further demonstrate the fuctional properties of personalized driver genes, we performe the enrichment analysis of the personalized driver genes in KEGG pathways to determine if the personalized driver genes are enriched in certain biological pathways.The enrichment results (including the pathway name, patient-occurred frequency and related combinational drugs) are shown in **Table S3**, from which we can find that 70% of these identified biological pathways are related with breast cancer, 52.5% related with lung cancer. These results indicate that CPGD can effectively identify the cancer-related pathways which are potentially targeted by drugs. Interestingly, we also find that most of the reported breast cancer-related pathways are enriched in many patients’ data with high frequency (>0.6), and some pathways with low frequency (<0.3) such as cell cycle have still been reported as targeted pathways for cancer therapy in previous studies^39^.

### Risk assessment by drug pairs with co-targeting on personalized driver genes

Recently some researchers developed a network-based methodology to identify anti-cancer drug pairs by exploring the “drug-drug-disease” combinations in the human interactome^9^. They found that anti-cancer drug pairs have a Complementary Exposure to the disease gene module,that is, both of the pairise combinational drugs can hit the targets overlapping with disease module, but two drugs of the pairise combinational drugs have separate (non-overlapping) targets^9^. Holding the similar assumption, we aim to uncover the efficacious anti-cancer combinational drugs according to the personalized driver genes identified with CPGD on BRCA cancer.

To identify anti-cancer drug pairs for risk assessment of BRCA cancer, we first collecte the interactions between drugs and genes, as well as the disease gene module of breast cancer. The interactions between drugs and genes are extracted from the combinatorial drugs and gene interaction network. The breast cancer-related genes were collected by Quan et al.^32^ from the Unified Medical Language System (UMLS)^40,41^. A total of 2341 breast cancer related genes is provided in **Table S4**. We select top 10 candidate combinational drugs for each individual cancer patient, identifying 12 drug pairs that have Complementary Exposure. This computational subprocedure can be seen in **Figure S2** (Additional file1). **Table 1** lists the name of 12 drug pairs, targeting personalized driver gene and patient frequency. Based on targeting personalized driver genes of these drug pairs, we carry out the risk assessment of these 12 drug pairs by using *SurvExpress* tool ^26^on TCGA BRCA data which contains 962 patients.

**Table 1.** Information of anti-cancer targeting driver genes and drug pairs for breast cancer

| Drug Pair ID | Drug A | Targeting driver genes of drug A | Drug B | Targeting driver genes of drug B | Patient frequency |
| --- | --- | --- | --- | --- | --- |
| DP-1 | DOXORU<br>BICIN | CYP1B1,CYP2C9,CYP3A4,<br>CYP3A5,MAP2,MAP4,MAP<br>T,NR1I2,RRM1,TUBB1,CY<br>P2C19 | PACLITAX<br>EL | CBR1,EGFR,ERBB2,RRM2 | 0.2738 |
| DP-2 | DOXORU<br>BICIN | CYP1B1,CYP2C9,CYP3A4,<br>CYP3A5,MAP2,MAP4,MAP<br>T,NR1I2,RRM1,TUBB1,CY<br>P2B6,CYP2C19 | GEMCITA<br>BINE | ABCB1,WRAP53,BIRC5 | 0.9910 |
| DP-3 | DOXORU<br>BICIN | C1S,CYP1B1,CYP2C9,CYP<br>3A4,CYP3A5,KDR,MAP2,<br>MAP4,MAPT,NR1I2,TUBB<br>1,CYP2B6,CYP2C19 | NERATINI<br>B | ABCB1,EGFR,ERBB2,TP53<br>,WRAP53 | 1 |
| DP-4 | DOXORU<br>BICIN | C1S,CYP1B1,CYP2C9,CYP<br>3A4,CYP3A5,MAP2,MAP4,<br>MAPT,NR1I2,TUBB1,CYP2<br>B6,CYP2C19 | LAPATINI<br>B | EGFR,ERBB2,TP53,WRAP<br>53 | 1 |
| DP-5 | PACLITA<br>XEL | C1S,CBR1,EGFR,ERBB2,C<br>YP2B6 | TRASTUZ<br>UMAB | ABCB1,ABCC1,ABCG2,C<br>YP1B1,CYP2C9,CYP3A4,C<br>YP3A5,MAP2,MAP4,MAP<br>T,NR1I2,TP53,TUBB1,WR<br>AP53,ABCC2 | 1 |
| DP-6 | GEMCIT<br>ABINE | ABCB1,WRAP53,BIRC5 | PACLITAX<br>EL | CBR1,CYP2B6,RRM2 | 0.8303 |
| DP-7 | CYCLOP<br>HOSPHA<br>MIDE | ATP7A,CYP3A4,CYP3A5,E<br>GFR,MAP2,MAP4,NR1I2,S<br>LCO1B3,TUBB1,MAPT | LAPATINI<br>B | PIK3CA,WRAP53 | 0.0714 |
| DP-8 | DOCETA<br>XEL | CYP2B6,CYP2C19,CYP2C9<br>,CYP3A4,EGFR,ERBB2 | LAPATINI<br>B | PIK3CA,WRAP53 | 0.0714 |
| DP-9 | FLUORO<br>URACIL | FLT1,KDR,KIT,PDGFRB | SUNITINI<br>B | ABCB1,IGFBP3,SLC29A1,<br>TP53,TYMS,WRAP53 | 0.0178 |
| DP-10 | FLUORO<br>URACIL | FLT1,KDR,KIT,MAPK11,M<br>APK12,PDGFRB,MAPK8,M<br>APK10 | SORAFENI<br>B | ABCG2,CYP3A4,ERCC1,I<br>GFBP3,SLC29A1,TP53,TY<br>MS,WRAP53 | 0.0803 |
| DP-11 | CARBOP<br>LATIN | ABCG2,CBR1,EGFR,ERBB<br>2,PIK3CA,WRAP53,XRCC4 | DOXORUB<br>ICIN | ABCB1,BRCA1,CYP2C9,C<br>YP3A4,CYP3A5,MAP2,MA<br>P4,MAPT,NR1I2,SLCO1B3,<br>TUBB1,CYP2C19 | 0.0178 |
| DP-12 | CARBOP<br>LATIN | ATP7A,C1S,EGFR,ERBB2,<br>XRCC4 | TRASTUZ<br>UMAB | ABCB1,BRCA1,CYP3A4,C<br>YP3A5,MAP2,MAP4,MAP<br>T,NR1I2,SLCO1B3,TUBB1 | 0.0178 |

As is shown in **Figure 7**, we find that among these 12 drug pairs, 11 drug pairs can actually divide all patients into discriminative high-risk and low-risk groups (p-value<0.05). From the results in **Figure 8** and **Figure S4-S6** (**Additional file1**), we can also find that among the 12 drug pairs, the *p*-value of 8 drug pairs are less than that of each single drug, indicating that the combined therapeutic effect of these 8 drug pairs are better than a single drug.

**Fig.7.**
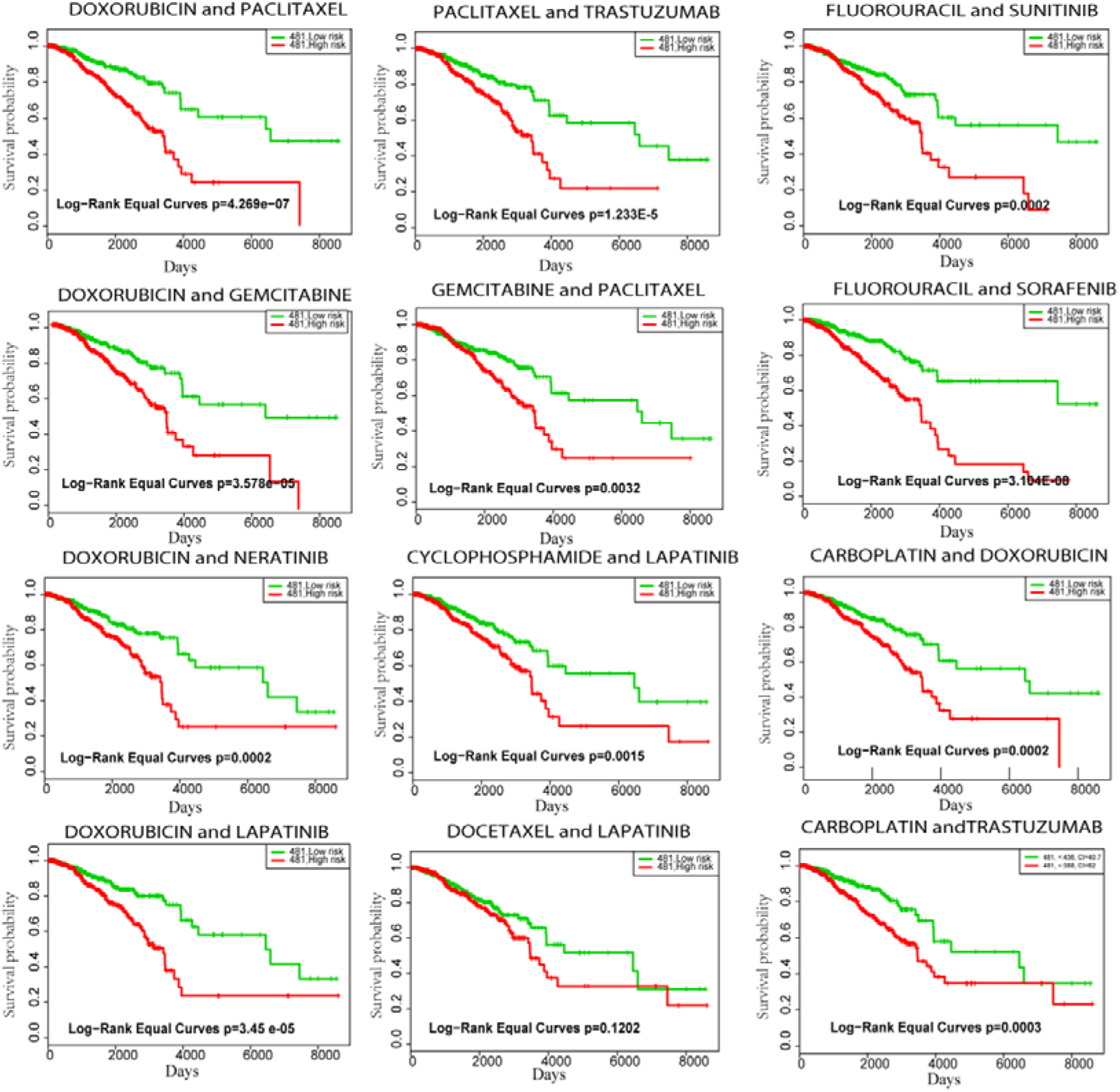
Results of the risk assessment of anti-cancer drug pairs for BRCA cancer.

**Fig.8.**
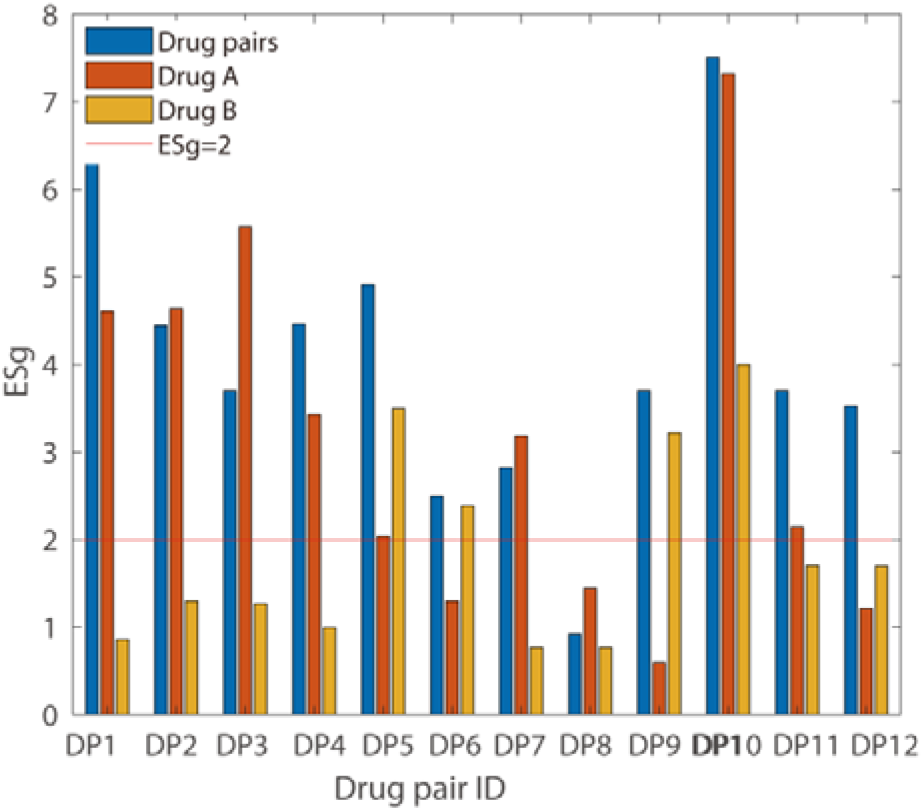
The p-value of combined drugs therapy and single drug therapy on BRCA cancer dataset. ESg=−log_10_(p-value).

### Cancer subtyping by quantifying side effect of drug pairs

To precisely quantify the side effect of drug pairs for mining cancer subtypes, we first obtained the personalized driver genes within two reliable cancer driver genes sets (i.e., the cancer census gene set^42^ and Network Cancer gene set^43^), then calculated the *side effect score* of a given drug pair by quantifing their side effect on personalized driver genes within cancer driver genes set for each patient. For an individual patient, the computational procedure of calculating the side effect score for a given drug pair is shown in **Figure S3** (Additional file1). Finally, we got the number of drug pairs with an *aggravating effect (side effect* score >0), and the number of drug pairs with *enhancing effect* (side effect score <0), which are used as two features of side effect signatures of individual patients.

By considering the side effect signatures of individual patients, we find that the patients in the breast cancer dataset can be signifacantly classified into three distinct subtypes (Figure 9A). Furthermore, we also evaluated the biological characteristics of these three subtypes from the following respects: i) We took the fraction of patient deaths and the fraction of patients with later stages in clinical (stage III and stage IV) as the index of significant risk assessment of different subtpes (**Figure 9B**), respectively. From **Figure 9B**, we can see that the greater the value of *aggravating effect*, the higher the risk effect on a specific patient subtype. These results show that the side effect signatures of these 3 subtypes are strong correlated with the risk effect in clinical. ii) Furthermore, to test the effect of 12 drug pairs (in **Table 1**) on three subtypes, we take the fraction of subtype patients with high risk as an index of risk assessment of these drug pairs on specific subtype (**Figure 9C**). From **Figure 9C**, we can see that 100% of subtype 1 patients are in high-risk groups for drug pair 1 (DOXORUBICIN and PACLITAXEL), 2 (DOXORUBICIN and GEMCITABINE), 5 (PACLITAXEL and TRASTUZUMAB), 6 (GEMCITABINE and PACLITAXEL), 8(DOCETAXEL and LAPATINIB), and 11(CARBOPLATIN and DOXORUBICIN), while only 71% of subtype 1 patients are in high-risk groups for drug pair 10 (DOXORUBICIN and PACLITAXEL). These results show that the risk effects of different drug pairs on a specific subtype vary in combination therapy.

**Fig.9.**
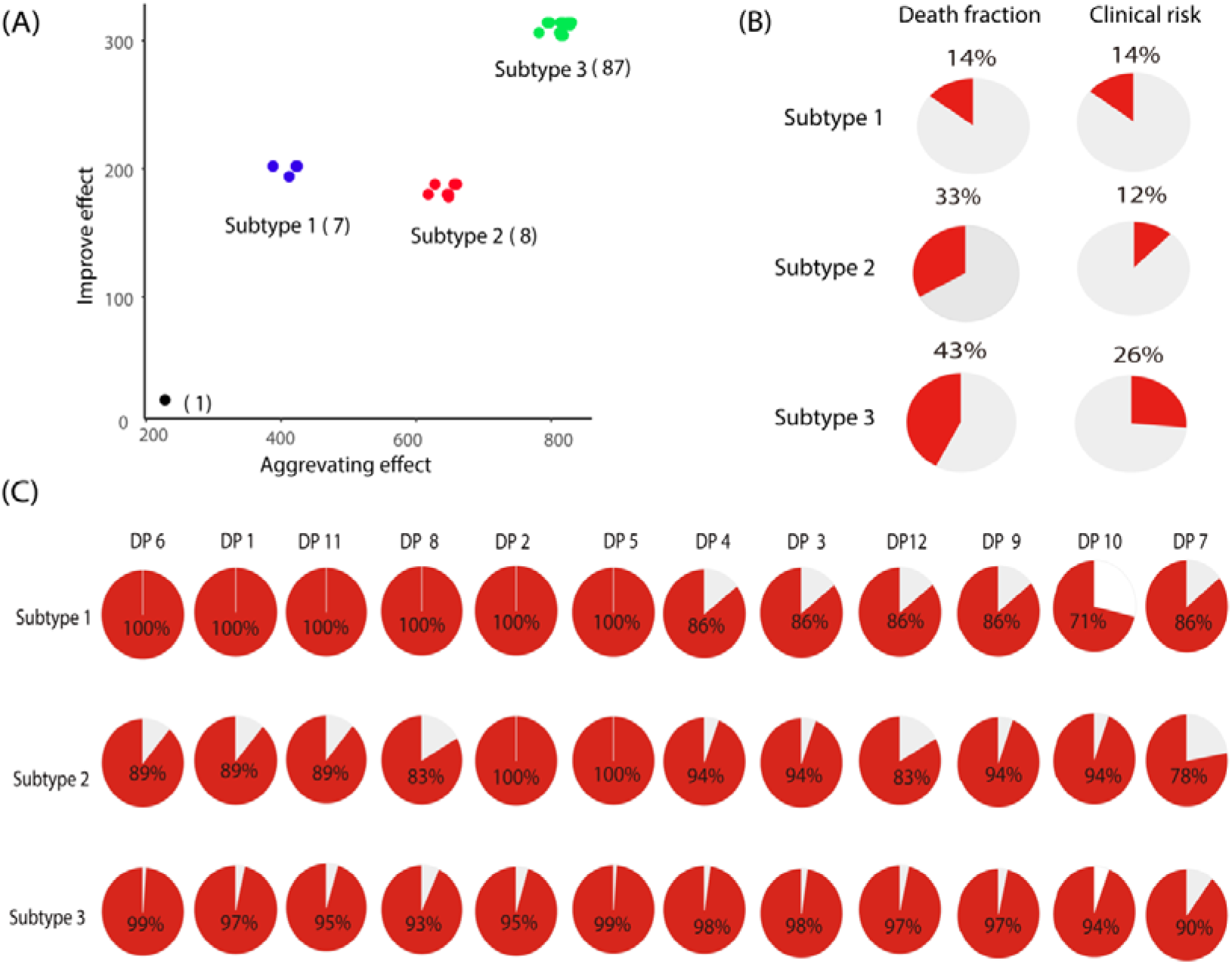
Results of the cancer subtyping generated by quantifying side effect of the drug pairs which target on the personalized driver genes. (**A**) The subtype classification of patients in the breast cancer dataset based on the number of drug pairs with an aggravating effect and an enhancing effect. (**B**) Results of the fraction of dead patients and the fraction of patients with stage III and stage IV based on our identified subtypes and clinically subtypes. (**C**) The risk assessment of 12 drug pairs on three subtypes.

## Discussions

In an effort to combat drug resistance in targeted therapy, combinatorial drugs are increasingly being used as the care treatment strategies for cancers^44^. Current methods of identifying combinational drugs can be categorized into: i) Network-based combinational drugs identification methods with *a priori* knowledge of gene regulatory networks^45–47^; ii) Machine learning-based combinational drugs identification methods, such as logistic regression^48^, random forest^49^ and deep learning^50^. However, current existing methods do not consider the personalized sample information (e.g., the personalized omics data), and thus fail to identify patient-specific combinational drugs. Therefore, it is a critical challenge how to develop effectively computational methods for discovering efficacious combinatorial drugs from the analysis of large biological and biomedical datasets.

Recently, many studies have demonstrated that targeted driver genes (i.e., candidate drug targets) can provide critical information for drug discovery and drug repurposing^11^. Coincidentally, structure network controllability methods mainly focus on how to identify driver nodes to determine the state transition of the whole (biological) network from the initial state to the desired state. For example, Single sample Controller Strategy (SCS) ^16^ and Personlaized Network Control model (PNC)^25^ have applied the structure network controllability to a single patient system to find personalized driver genes. However, there is currently no feasible way of distilling these structure network controllability methods into discovery of personalized combinatorial drugs.

In this work, we consider the personalized drug combination identification as a typical structure network controllability problem, then present a novel method (namely CPGD) to identify the target genes and further screen the combination drugs. CPGD first uses a new Paired-SSN method to construct the personalized transition network, then introduces the Weight-NCUA (a kind of strucural network controllability method) to find the optimal personalized driver genes. Technically, Weight-NCUA considers how to push a individual system of cancer from one attractor (tumor) to another attractor (normal, or approximate normal) through perturbing optimal personalized driver genes by combinatorial drug activation. From structure network controllability perspective, the optimal personalized driver genes theoretically can determine the state transition between disease state and normal state (or approximate normal state). We evaluate the CPGD performance from the following aspects: i) prioritizing personalized combinatorial drugs to evaluate the ability of predicting clinical efficacious combinational drugs, ii) exploring risk assessment of paiwise combinatorial drugs, and iii) enhancing cancer subtyping by side effect quantification on personalized driver genes.

The results BRCA and LUSC datasets show that our CPGD is better than other existing methods in terms of *Precision*. The drug pairs identified with CPGD can divide patients into discriminative high-risk and low-risk groups for effective risk asessment. In addition, CPGD can effectively mine the cancer subtypes on breast cancer dataset by quantificating the side effect of combinatorial drugs on co-targeting personalized driver genes for individual patients.

Overall, the results suggest that the discovery of personalized combinational drugs in cancer could benifit from network controllability-based approaches. However, this study doesn’t consider the dynamic change of drug concentration, which may lead to some false postives for the identification of efficacious combinational drugs. In addition, biological experimental validation and prospective clinical trials should be conducted to verify the discovery of CPGD. A more complete, systematic gene interaction network and drug-target network may further improve the performance of the network controllability-based approaches.

## Materials and Methods

### Combinatorial drugs-genes interaction networks

The interactions between combinatorial drugs and genes were collected by Quan et al. ^32^ from Drug Combination Database (DCDB)^51^, drug-gene Interaction database (DGIdb)^52^, DrugBank^53^ and Therapeutic Target Database (TTD)^54^. The combinatorial drugs-genes interaction network contains 342 combinatorial drugs and 5,788 interaction edges (listed in **Table S1**). It has been reported that 122 of 342 combinatorial drugs (35.7%) are efficacious for treating cancer, and this ratio is significantly higher than the background ratio of anti-cancer combinatorial drugs (14.8%) in DCDB ^32^. Therefore, these interactions are efficacious sources for the combinatorial drugs identification. Among these, 32 and 17 combinatorial drugs (listed in **Table S2**) derived from DCDB are efficacious in treating breast cancer and lung cancer, respectively.

### Human drug-target network with activation and inhibition interactions

To analyze the risk assessment of cancer patients, the human drug-target network with activation and inhibition interactions information is added to the network-based investigation of drug-target interactions^55^.

### Paired-SSN method

For Paired-SSN method^25^, the co-expression networks of the tumor sample and normal sample for an individual patient are separately built by using SSN method^56^. For one single sample, the significant Z-score of an edge between gene *i* and *j* are calculated by using the following formulars:

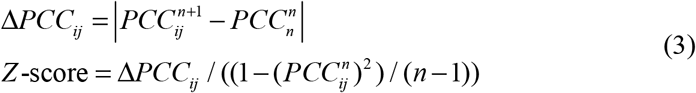

where 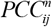 is the Pearson correlation coefficient (PCC) of an edge between gene *i* and *j* in the reference network with *n* reference samples (here, refenence samples are all normal samples for specific cancer); 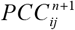 is the Pearson correlation coefficient of the edge between gene *i* and *j* in the perturbed network with one additional sample, that is, adding a new sample (normal sample or tumor sample) for an individual patient to the reference group samples. The P-value of an edge could be obtained from the statistical Z-score. All of the interacted edges with significant differential correlations (e.g., P-value<0.05) are adopted to build SSNs for the normal and tumor samples (**Figure 1**), that is, the edges of a single sample network (for normal sample or tumor sample) exist in both gene interaction network and the co-expression network.

Then, the personalized personalized gene interaction networks are constructed by using the following criterion. If the P-value of the edge between gene *i* and gene *j* is less than 0.05 in the tumor sample network but greater than 0.05 in the normal sample network, or greater than 0.05 in the tumor sample network and less than 0.05 in the normal sample network, the edge between gene *i* and gene *j* is retained to constitute the personalized personalized gene interaction network. Additionally, we calculate the edge weight *e_ij_* between gene *i* and gene *j* with the following formula:

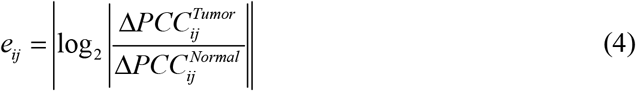

To quantify the regulatory score of each gene on the personalized personalized gene interaction network, the node weight *W_i_* is calculated with the following formula:

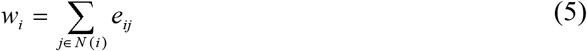

where *N*(*i*) represents the neighboring node set of node *i* in the personalized personalized gene interaction network. Therefore, the personalized personalized gene interaction networks can reveal the significant gene interactions between the normal and tumor samples for each patient during cancer development.

### Weight-NCUA method

Theoretically, Weight-NCUA uses the following equation to represent the dynamic behavior of a personalized personalized gene interaction network:

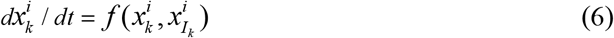

where 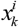 denotes the expression state of a gene *k* for the patient *i, I_k_* represents the neighborhood gene set of gene *k*, and *f* represents the dynamic behavior control law of the personalized personalized gene interaction network for patient *i*, satisfying the continuous differentiability, dissipativity, and decay conditions^57,58^.

This dynamic Eq.6 represents the dynamic behavior of the gene expression level in the personalized gene interaction network. For a personalized gene interaction network, we assume that each edge is bi-directional. Thus, we convert it into a bipartite network in which the up nodes and bottom nodes are the nodes and the edges of the original network. If the node *v_i_* in up side is one of nodes for edge *V_j_* in the bottom side, then *V_i_* and *V_j_* are linked in the bipartite network. Based on the FVS structure controllability theory, Weight-NCUA selectes the dominanting nodes set *M* in the up side that cover the nodes in the bottom side as the driver nodes for determining the state of the network.

It is known that different dominanting nodes sets may generate multiple driver-node sets, resulting in a potential bottleneck for identifying the combinational drugs of the individual patients. Therefore, we introduce an index to indicate the quality of the selected personalized driver genes:

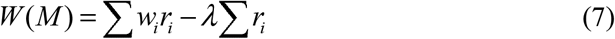

where *W_i_* denotes the confidence score of gene *i*; *r_i_* is an indicative variable, when gene *i* is selected as the personalized driver genes, *r_i_* = 1, otherwise *r_i_* = 0; Σ*W_i_r_i_* denotes the network scores of candidate gene sets; Σ*r_i_* denotes the number of candidate personalized driver genes; and *λ* is the balance parameter to adjust the network scores of candidate personalized driver genes and the number of candidate personalized driver genes.

For Weight-NCUA, we expect that the personalized driver genes not only contain the minimum number, but also have the maximum network scores. It is need to further measure the quality of candidate personalized driver genes set. Thus, we select the personalized driver genes by solving the following linear integer programming:

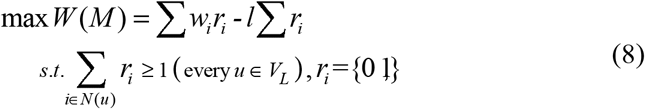

where *V_L_* denotes the bottom nodes in the bipartite graph and *N*(*u*) denotes the neighborhood nodes in the bipartite graph.

The objective function is to obtain the personalized driver genes with the minimum number but the maximum network scores. The restriction condition is to ensure that all the edges of the original network in bipartite network can be covered. Under the dynamic behavior, the state of all genes in a patient-specific network can be regulated by the personalized driver genes. Above optimization problem can be efficaciously solved by using the LP-based classic branch and bound method^59,60^.

### Calculating the side effect scores for a given drug pair

To calculate the *side effect score* of a given drug pair, we first collect the drug-target network with the information of activation and inhibition interaction. Then we give the classification results of sharing targets of two drugs for characterizing the effect of a given drug pair. The configurations (i.e, (+,+) and (−,−)) of the sharing targets of two drugs are referred as coherent, where the action of one drug on the sharing targets is reinforced by the presence of a second drug. The configuration (+,−) of the sharing targets of two drugs is called incoherent, where the action of one drug on the sharing targets is mitigated by the presence of a second drug. Finally, the *side effect score* can be calculated by using the following formula^55^:

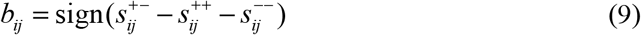

where 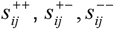 respectively denote the number of sharing targets with configurations (+, +), (+, −) and (−, −) for drug pair (*i, j*).

### Enrichment pathway analyzing of the personalized driver genes in KEGG pathway dataset

To identify the enrichment pathways of the personalized driver genes, we compute the *p*-value of enrichment pathways by using the hyper-geometric test^61^,

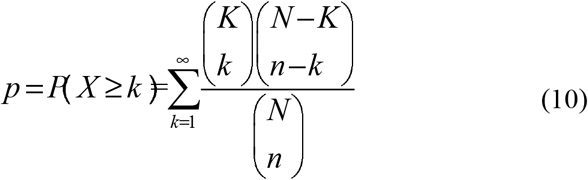

where *N* is the number of genes in the gene interaction network, *n* is number of the personalized driver genes, *K* is the number of genes in a given pathway from the KEGG pathway dataset, and *k* is the number of the personalized driver genes within the given pathway. If p-value for a given pathway is less than 0.05, the personalized driver genes are significantly enriched in the given pathway.

### Statistical test of the functional and structural properties for the personalized driver genes

#### Mutation profile test

We use the following equation to evaluate whether the personalized driver genes are significantly enriched in the set of mutated genes for an individual patient.

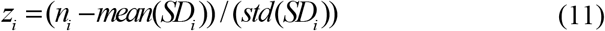

where *n_i_* is the the number of intersected gene between mutated genes and personalized driver genes for patient *i. SD_i_* is the distribution of the number of intersected gene between mutated genes and random generated genes. The number of randomly generated genes is equal to that of personalized driver genes. Mean and standard deviation of *SD_i_* are calculated from 100 simulations of random datasets. Based on the z-score, we can obtain the empirical P-values for each patient. The alternative hypothesis (H0) is *n_i_* = *n*_0_, and the alternative hypothesis (H1) is *n_i_* > *n*_0_, where *n*_0_ is mean value of *SD_i_* in the mutation profile test.

#### Epistasis-interaction test

Some studies show that combinatorial drugs aiming at the epistatic genes may have higher medicinal potential^11^. The epistatic genes are defined as the genes that are interacted each other in the gene interaction network. For the epistasis interaction test, we first calculate the number of edges among the personalized driver genes, then generate 100 random networks by using randomization-based test^62^, each of which maintains the topological characteristics of the original network (i.e., node degree), recalculating the number of edges among the personalized driver genes. Finally, we use Eq. 11 to get the *P*-value for evaluating whether the personalized driver genes are significantly more connected in the gene interaction network than the random networks. In the calculation of *z*-score, *n_i_* is the the number of edges among the personalized driver genes for patient *i, SD_i_* is the distribution of the number of edges among personalized driver genes in the random networks. Mean and standard deviation of *SD_i_* are calculated from 100 simulations. For the epistasis-interaction test, the alternative hypothesis (H0) is *n_i_* = *n*_0_ and the alternative hypothesis (H1) is *n_i_* > *n*_0_, where *n*_0_ is mean value of *SD_i_* in the epistasis interaction test.

## Supporting information

supplemental files including addrtional file and Tabes S1,S2,S3 and S4

## Code availability

The implementation of our CPGD in MATLAB can be freely downloaded from https://github.com/NWPU-903PR/CPGD.

## Data availability

The gene expression data, combinatorial drugs-gene interaction data and drug-gene interaction data with activation and inhibition interactions can be freely downloaded from https://github.com/NWPU-903PR/CPGD.

## Author contributions

WFG developed the method and executed the experiments. WFG and TZ made analysis and wrote this paper. WFG, SWZ, TZ and LNC revised the manuscript. SWZ and LNC supervised the work, made critical revisions of the paper, and approved the submission of the manuscript. All authors read and approved the final manuscript.

## Competing interests

The authors have declared no competing interests.

## Acknowledgements

We thank Professor Jianxi Gao from Rensselaer Polytechnic Institute, U.S.A., for giving us useful and valuable comments. We also thank Dr. Meng-Yuan Liu for providing us the combinational drugs and gene interaction data, synthetic lethality genes interactions data.

## Fundings

This paper was supported by the National Natural Science Foundation of China (61873202, 61473232, 91430111, 31930022, 31771476, 81471047 and 11871456), National Key R&D Program (2016YFC0903400, 2017YFA0505500), Shanghai Municipal Science and Technology Major Project (No. 2017SHZDZX01), and Natural Science Foundation of Shanghai (17ZR1446100).

## Notes

#### Summary of Updates

We have reorganized our manuscript which improves the orignal version.

