## supplemental files including addrtional file and Tabes S1,S2,S3 and S4 for "A novel network controllability algorithm to target personalized driver genes for discovering combinational drugs of individual cancer patient": Additional_file1_Supplementary_manuscript.docx

**Supplementary note 1: supplementary figures and supplementary tables**

Figure S1. The comparisons of number of driver nodes for Weight-NCUA (the second step of PDC) and other structure network control methods on BRCA (A) and LUSC (B).

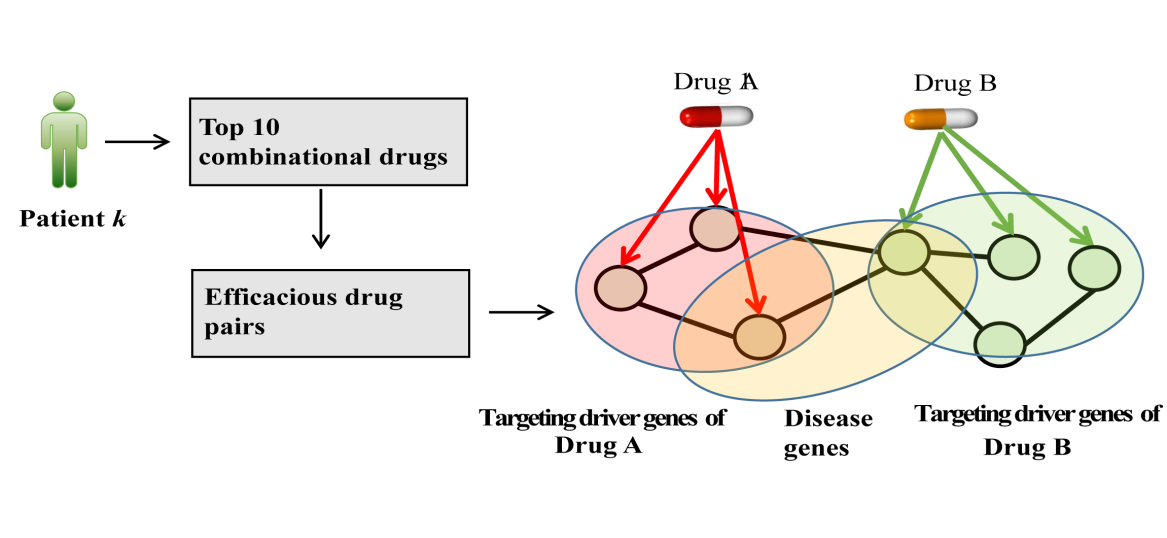

**Figure S2 The computational procedure for** exploring risk assessment of **anti-cancer drug pairs.** To explore risk assessment of pairwise drug combinations, we firstly selected top 10 candidate combinational drugs for each patient. Then among these candidate combinational drugs, we selected drug pairs that have Complementary Exposure. e.g., drug pairs can both hit the personalized driver genes overlapping with disease module but have separate (non-overlapping) targeted personalized driver genes 9. The interactions between drugs and personalized driver genes were extracted from the drug combinations and gene interaction network (Materials and methods) while the cancer-related genes can be identified by using the Unified Medical Language System (UMLS) ^24-26^. Finally we selected efficacious drug pairs with significant survival analysis results (p-value<0.05) for risk assessment from drug pairs that have Complementary Exposure by using their targeting personalized driver genes as the input of SurvExpress tool ^27^ .

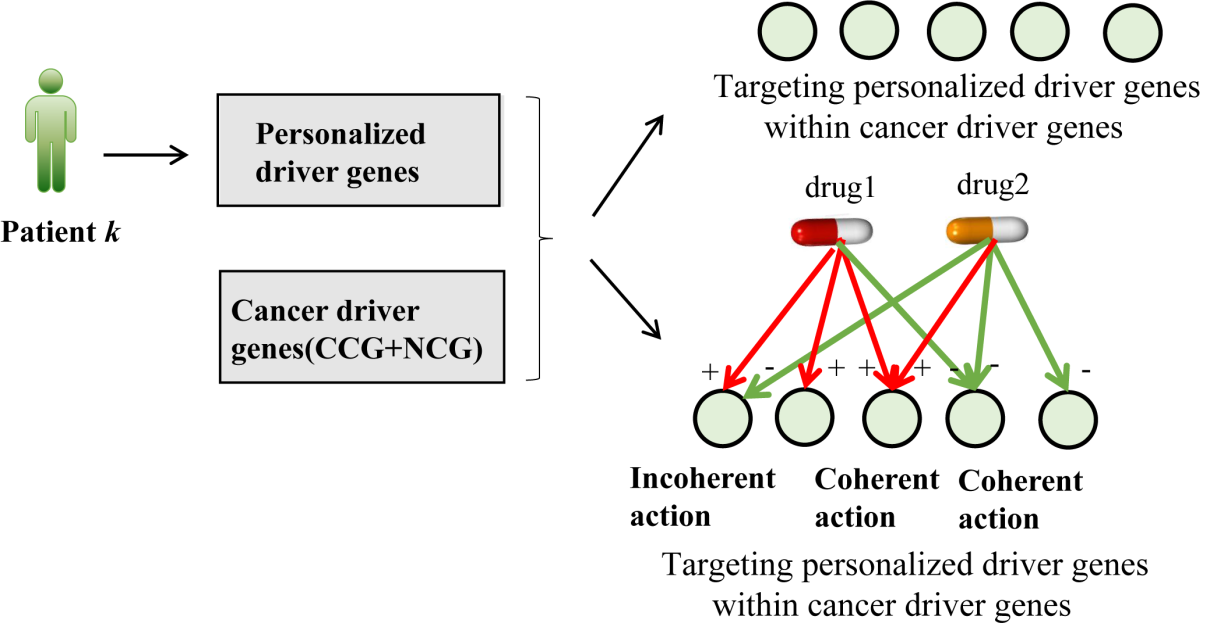

**Figure S3 The computational procedure for calculating the side effect score of a given drug pair for each patient.** We calculated the side effect score of a given drug pair by quantifying side effect on personalized driver genes in the drug-target network with positive and negative interactions for each patient. When two drugs act simultaneously on the same target, their action of two combinations (+,+) and (−,−) will be referred to as *coherent* action and the action of two combinations (+,−) is called incoherent action. By attaching signs to the mechanisms of action, the *side effect score* can be calculated by using
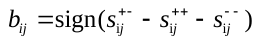
 where *s_ij_* is the number of common targets of the drug pair (*i*, *j*) in the drug-target network with positive and negative interactions for each patient (Materials and Methods).

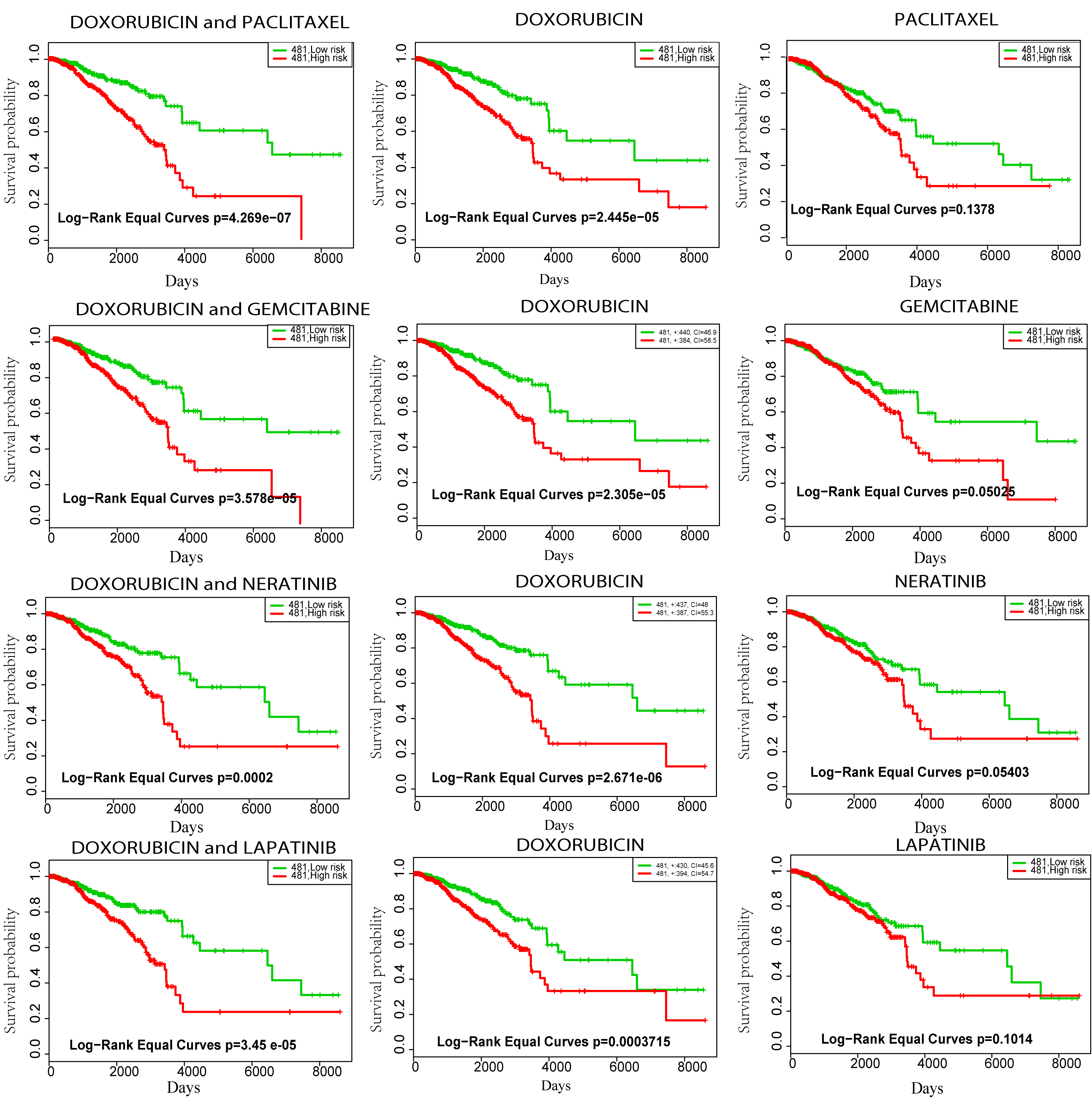

**Figure S4** The p-value of drug pairs and each single drug for drug pairs 1-4 on BRCA cancer data set. The information of the drug pairs name and targeting personalized driver gene is shown in Table 1 of manuscript.

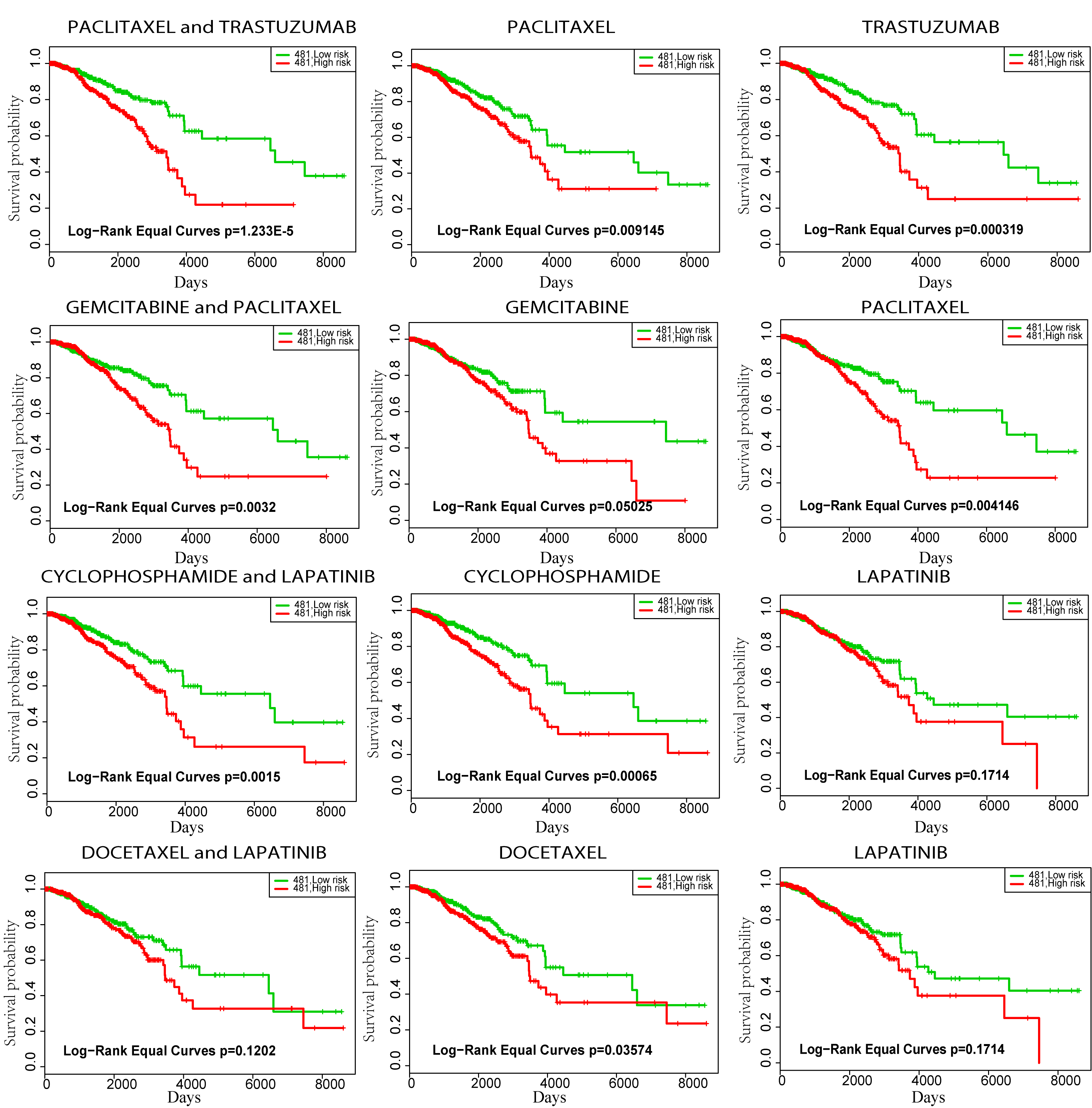

**Figure S5** The p-value of drug pairs and each single drug for drug pairs 5-8 on BRCA cancer data set. The information of the drug pairs name and targeting personalized driver gene is shown in Table 1 of manuscript.

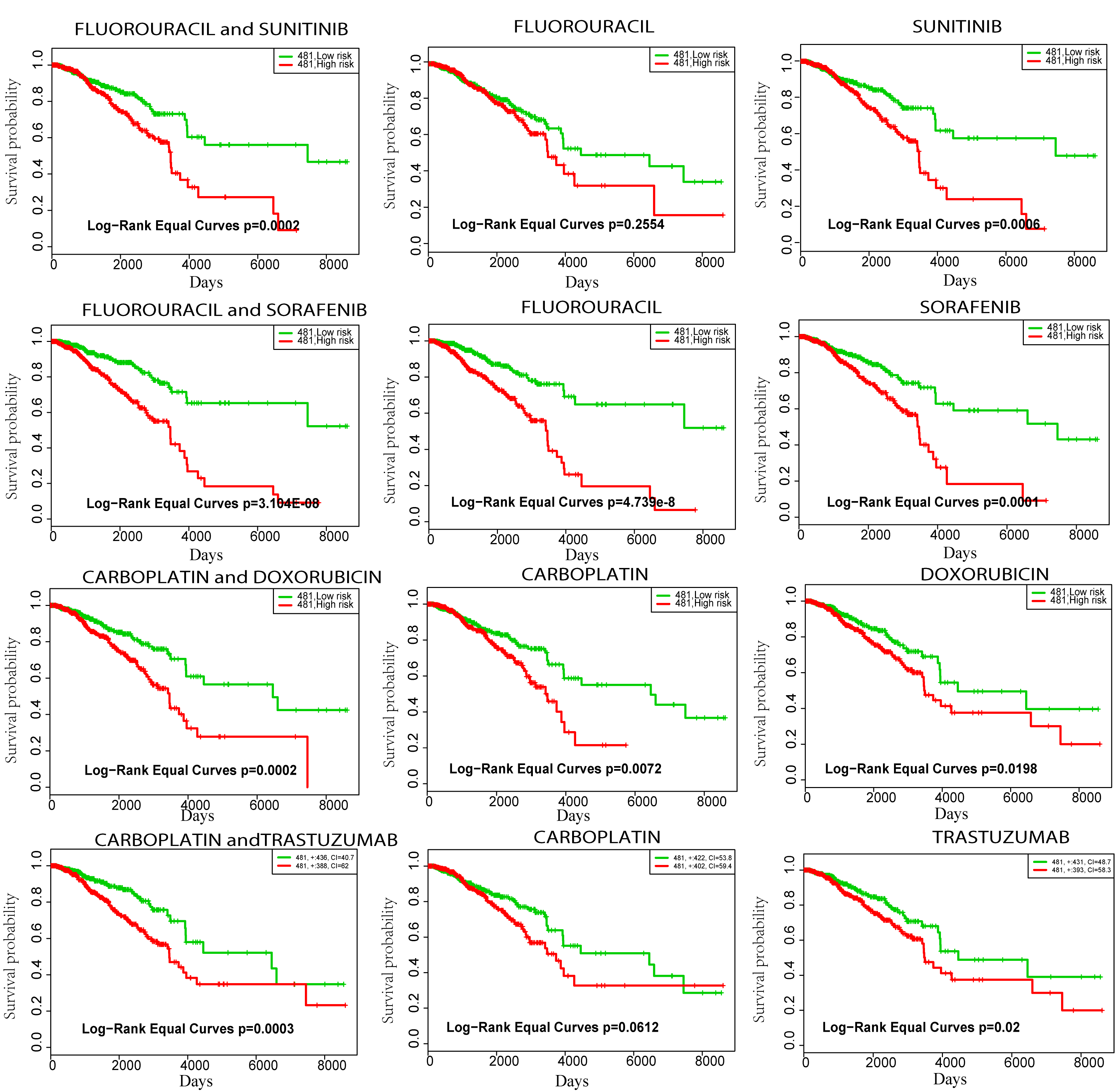

**Figure S6** The p-value of drug pairs and each single drug for drug pairs 9-12 on BRCA cancer data set. The information of the drug pairs name and targeting personalized driver gene is shown in Table 1 of manuscript.
